# Novel and potent inhibitors targeting DHODH, a rate-limiting enzyme in *de novo* pyrimidine biosynthesis, are broad-spectrum antiviral against RNA viruses including newly emerged coronavirus SARS-CoV-2

**DOI:** 10.1101/2020.03.11.983056

**Authors:** Rui Xiong, Leike Zhang, Shiliang Li, Yuan Sun, Minyi Ding, Yong Wang, Yongliang Zhao, Yan Wu, Weijuan Shang, Xiaming Jiang, Jiwei Shan, Zihao Shen, Yi Tong, Liuxin Xu, Chen Yu, Yingle Liu, Gang Zou, Dimitri Lavillete, Zhenjiang Zhao, Rui Wang, Lili Zhu, Gengfu Xiao, Ke Lan, Honglin Li, Ke Xu

**Affiliations:** State Key Laboratory of Virology, College of Life Sciences, Wuhan University, Wuhan 430072, China; Shanghai Key Laboratory of New Drug Design, State Key Laboratory of Bioreactor Engineering, School of Pharmacy, East China University of Science and Technology, Shanghai 200237, China; State Key Laboratory of Virology, Wuhan Institute of Virology, Center for Biosafety Mega-Science, Chinese Academy of Sciences, Wuhan 430071, China; CAS Key Laboratory of Molecular Virology and Immunology, Institut Pasteur of Shanghai, Chinese Academy of Sciences; University of Chinese Academy of Sciences, 320 YueYang Road, Shanghai 200031, China

## Abstract

Emerging and re-emerging RNA viruses occasionally cause epidemics and pandemics worldwide, such as the on-going outbreak of coronavirus SARS-CoV-2. Existing direct-acting antiviral (DAA) drugs cannot be applied immediately to new viruses because of virus-specificity, and the development of new DAA drugs from the beginning is not timely for outbreaks. Thus, host-targeting antiviral (HTA) drugs have many advantages to fight against a broad spectrum of viruses, by blocking the viral replication and overcoming the potential viral mutagenesis simultaneously. Herein, we identified two potent inhibitors of DHODH, S312 and S416, with favorable drug-like and pharmacokinetic profiles, which all showed broad-spectrum antiviral effects against various RNA viruses, including influenza A virus (H1N1, H3N2, H9N2), Zika virus, Ebola virus, and particularly against the recent novel coronavirus SARS-CoV-2. Our results are the first to validate that DHODH is an attractive host target through high antiviral efficacy *in vivo* and low virus replication in DHODH knocking-out cells. We also proposed the drug combination of DAA and HTA was a promising strategy for anti-virus treatment and proved that S312 showed more advantageous than Oseltamivir to treat advanced influenza diseases in severely infected animals. Notably, S416 is reported to be the most potent inhibitor with an EC50 of 17nM and SI value >5882 in SARS-CoV-2-infected cells so far. This work demonstrates that both our self-designed candidates and old drugs (Leflunomide/Teriflunomide) with dual actions of antiviral and immuno-repression may have clinical potentials not only to influenza but also to COVID-19 circulating worldwide, no matter such viruses mutate or not.

## Introduction

Acute viral infections, such as influenza virus, SARS-CoV, MERS-CoV, Ebola virus, Zika virus, and the very recent SARS-CoV-2 are an increasing and probably lasting global threat^1^. Broad-spectrum antivirals (BSA) are clinically needed for the effective control of emerging and re-emerging viral infectious diseases. However, although great efforts have been made by the research community to discover therapeutic antiviral agents for coping with such emergencies, specific and effective drugs or vaccines with low toxicity have been rarely reported^2^. Thus, unfortunately, there is still no effective drugs for the infection of the novel coronavirus SARS-CoV-2 at present, which outbreak in December 2019 firstly identified by several Chinese groups^3-5^, and now has quickly spread throughout China and to more than 90 other countries, infecting 101,923 patients and killing 3486 ones by March 7, 2020^6^.

Discovery of nucleoside or nucleotide analogs and host-targeting antivirals (HTAs) are two main strategies for developing BSA^7-9^. With the former drug class usually causing drug resistance and toxicity, the discovery of HTAs has attracted much attention^10^. Several independent studies searching for HTAs collectively end up to compounds targeting the host’s pyrimidine synthesis pathway to inhibit virus infections, which indicates that the replication of viruses is widely dependent on the host pyrimidine synthesis^11-25^. However, most of these compounds lack verified drug targets making subsequent drug optimization and further application impossible^11,13-15,17-19,21,25^. There are only a few inhibitors against pyrimidine synthesis that can be carried forward to animal studies, however, their antiviral efficacies were unsatisfactory or even ineffective at all^12,16,17,21,23-25^. For example, a pyrimidine synthesis inhibitor FA-613 without a specific target protected only 30.7% of mice from lethal influenza A virus infection when compared to the DAA drug Zanamivir (100%) in parallel^23^. Another two compounds, Cmp1^24^ and FK778^25^, which target DHODH, a rate-limiting enzyme in the fourth step of the *de novo* pyrimidine synthesis pathway, could only inhibit the DNA virus (CMV) replication in RAG^-/-^ mice, but their therapeutic effects on the upcoming diseases were unexplored. Therefore, more potent pyrimidine synthesis inhibitors, especially ones with the specific drug target, are urgent to be developed to prove whether such an HTA drug is valuable towards clinical use or has any advantages over DAA drugs in antiviral treatment.

To identify potent and low-toxicity DHODH inhibitors (DHODHi), we previously conducted a hierarchal structure-based virtual screening (**Fig. 1A**) against ∼280,000 compounds library towards the ubiquinone-binding site of DHODH^26^. We finally obtained two highly potent DHODHi S312 and S416 with IC_50s_ of 29.2 nM and 7.5 nM through structural optimization^27,28^, which are > 10-folds potent than the FDA approved DHODHi Teriflunomide (IC_50_ of 307.1 nM). By using these two potent inhibitors, we could fully evaluate DHODH as a valuable host target both in infected cells and *in vivo* in infected animals. We identified that targeting DHODH offers broad-spectrum antiviral efficacies against various RNA viruses, including the DAA-resistant influenza virus and the newly emerged coronavirus SARS-CoV-2. Especially, our potent DHODHi can protect 100% mice from lethal influenza challenge, which is as good as the DAA drugs, and is even effective in the late phase of infection when DAA drug is no longer responding.

**Figure 1.**
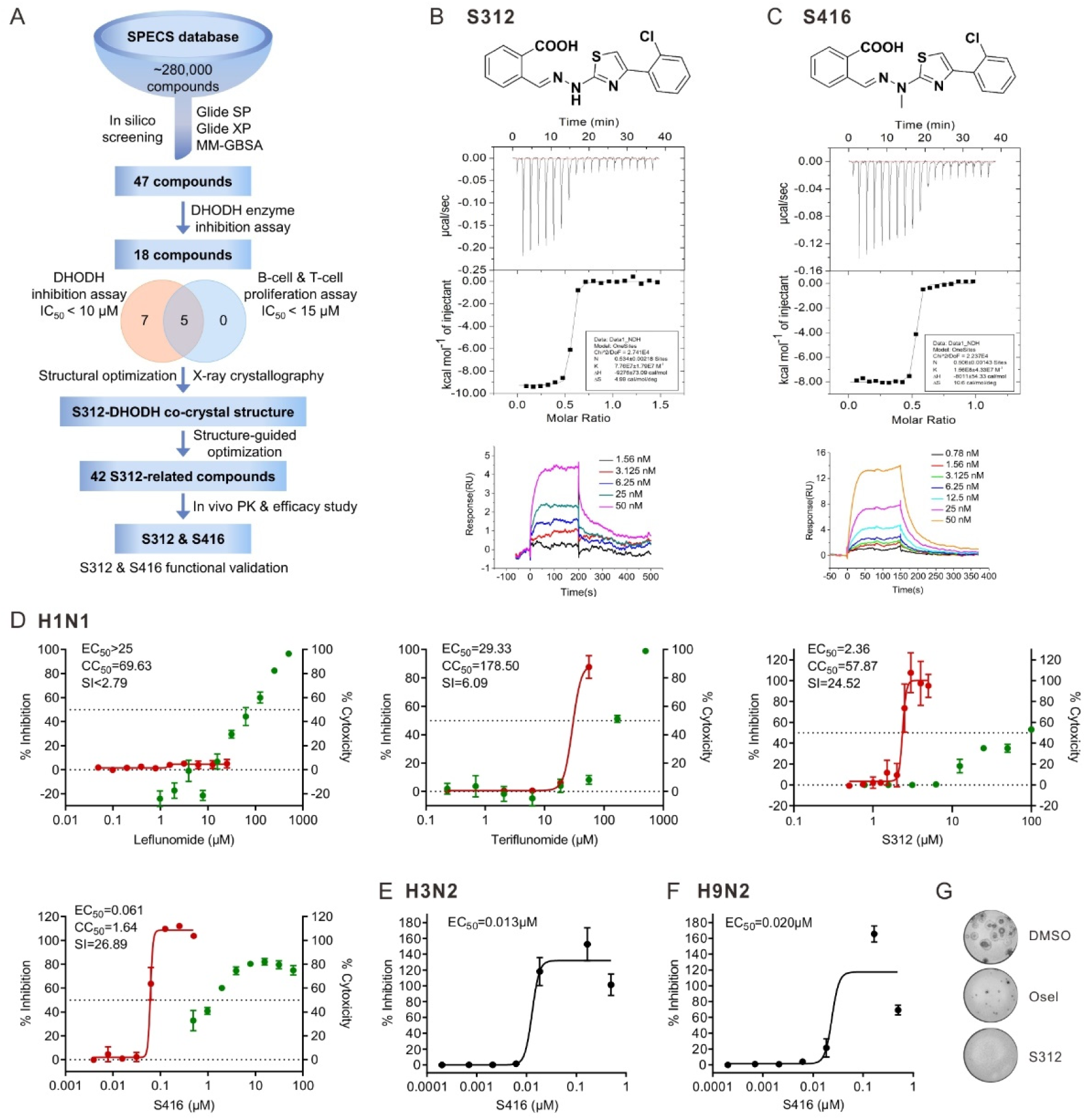
Discovery of potent DHODH inhibitors and their anti-influenza-A-virus activities. **(A)** The discovery and design diagram. Binding analysis of S312 **(B)** and S416 **(C)**. Thermodynamic analysis of the binding of S312 and S416 to DHODH were carried out at 25 °C on a MicroCal iTC200 instrument. Kinetic analysis of the binding of S312 and S416 to DHODH were performed with a Biacore T200 instrument. **(D)** Inhibitory activities of DHODH inhibitors against influenza A virus (A/WSN/33 [H1N1]). MDCK cells were infected with WSN virus (20 TCID_50_/mL) in the presence of increasing concentrations of compounds (Teriflunomide, Leflunomide, S312 and S416) for 72h. Inhibition efficiencies of these four compounds against the WSN virus (left-hand scale, red curve) and their cytotoxicity (right-hand scale, green curve) were all determined using cell viability assay. **(E) and (F)** Antiviral activities of S416 against influenza A virus H3N2 and H9N2. The experimental procedure and the detection method were the same as shown in (D). EC_50_ of the S416 was shown. The results (D, E, and F) are presented as a mean of at least three replicates ± SD. **(G)** Observation of virus morphology in the presence of the indicated compound. MDCK cells were treated with 5IC_50_ of S312 or Oseltamivir at the meantime of infection. Cells were fixed and stained after 48 h.p.i., Upper well, S312 (12.5 μM); Centre well, Oseltamivir (3 μM); Bottom well, Control (DMSO).

## Results

### The discovery of potent DHODH inhibitors with low toxicity *in vivo* and high antiviral efficacies in influenza-A-virus infected cells

By determination of the X-ray crystal structure of DHODH in complex with S416 (**Supplementary Data Fig. 1A, Table S1**), we verified the binding mode of S416 at the ubiquinone-binding site of DHODH that similar to S312. The binding free energies for S312 and S416 were -45.06 and -46.74 kJ/mol, and the binding equilibrium dissociation constants (*K*_D_) were 20.3 and 1.69 nM, respectively (**Fig. 1B and 1C**). Additionally, the two inhibitors exhibited a clear trait of fast-associating (*k*_on_) and slow-dissociating (*k*_off_) inhibition (**Table S2**), providing themselves as ideal drug candidates with a high level of target occupancy. Moreover, S312 and S416 showed proper half-lives (8.20 and 9.12 h, respectively) (**Table S3**), which are much shorter than that of Teriflunomide and Leflunomide, indicating that they may have less possibility to bring side effects from drug accumulation in the body (**Supplementary Data Fig. 1B and 1C**).

To examine the antiviral activities of these DHODHi, we use the influenza A virus as a model virus. A labor stain of A/WSN/33(H1N1, WSN) with 20 TCID_50_ was applied to infect MDCK cells, and serial dilutions of drugs (DMSO as controls) were added at the same time when cells were infected. Drug efficacies were evaluated by quantification of cell viability in both infected and non-infected cells, and the half-maximal effective concentration (EC_50_) and the half-cytotoxic concentration (CC_50_) of the indicated drug were obtained accordingly. The selectivity index (SI) was calculated by CC_50_/EC_50_. As shown in **Fig. 1D**, the antiviral effect of Leflunomide is hardly detectable at the cell culture level (EC_50_>25 μ M). However, Teriflunomide, the active metabolite of Leflunomide, exhibited a clear antiviral effect against the WSN virus (EC_50_=29.33μM, CC_50_=178.50μM, SI=6.08). As compared to Teriflunomide, the potent DHODHi S312 is ∼12-fold stronger (EC_50_=2.36μM) and S416 is ∼480-fold stronger (EC_50_=0.061μM) than Teriflunomide in their antiviral efficacies. We also tested different influenza A virus subtypes of H3N2 and H9N2. The antiviral efficacies of DHODHi followed the same pattern as they were against H1N1, which is S416>S312>Teriflunomide>Leflunomide in viral inhibitory efficacies (summarized in **Table 1**). The drug effective curve of S416 to H3N2 (EC_50_=0.013μM) and H9N2 (EC_50_=0.020μM) is shown in **Fig. 1E and 1F**. When we compared the drug efficacy by virus plaque assay, the results in **Fig. 1G** showed that the positive control DAA drug Oseltamivir (Osel) could reduce the plaque size to needlepoint size. However, the virus plaque in equivalent S312-treatment was not observable at all indicating that S312 is more efficient in inhibiting virus replication than Osel.

**Table 1.** Broad-spectrum antiviral efficacies of DHODH inhibitors.

| Virus type | IC<sub>50</sub>(μM) (SI*<sup>1</sup>) | | | | | |
| --- | --- | --- | --- | --- | --- | --- |
|  | S312 | S416 | Teriflunomide | Leflunomide | Oseltamivir | Brequinar |
| A/WSN/33 (H1N1) | 2.36 (25) | 0.061 (27) | 29.33 (6) | >25.00(<2.7) | 7.68 (680) | 0.240 (11.9) |
| A/DongHu/06 (H3N2) | 8.43 (7) | 0.013 (126) | 2.73 (32) | N.T.* | N.T.* | 0.022 (130) |
| A/GuangZhou/99 (H9N2) | 13.17 (4) | 0.020 (82) | 3.36 (26) | N.T.* | N.T.* | 0.060 (48) |
| Zika virus | 2.29 (26) | 0.021 (1087) | 17.72 (3) | N.T.* | N.T.* | 0.268 (48) |
| Ebola-replicon | 14.96 (7.9) | 0.018 (4746) | 6.43 (21) | N.T.* | N.T.* | 0.102 (127) |

SI\* value was equal to CC50/IC50
N.T. (no test)

**Table 2.** Antiviral abilities of DHODH inhibitors against Osel-resistant strain.

| Virus type | IC<sub>50</sub>(μM) | |
| --- | --- | --- |
|  | S312 | Oseltamivir |
| NAmut<sup>H275Y</sup> | 5.967 | ~1046 |
| Wild type | 3.077 | 43.01 |

The results in all indicate that DHODHi, especially S312 and S416 exhibited direct antiviral activities to different subtypes of influenza A viruses by shutting off virus multiplication more thoroughly than Osel.

### The broad-spectrum antiviral activities of DHODHi to Zika, Ebola and newly emerged SARS-CoV-2 viruses in infected cells

As all actuate infectious viruses rely on cellular pyrimidine synthesis process to replicate, it is reasonable to speculate that DHODHi have broad-spectrum antiviral efficacies. We, therefore, tested several highly impacted acute infectious RNA viruses. All compounds of Teriflunomide, Brequinar, S312, and S416, showed inhibitory effects against Ebola virus (EBOV) mini-replicon, with EC_50_ of 6.43, 0.102, 14.96 and 0.018μM, respectively (**Fig. 2A**). To our supersize, S416 showing relatively high cytotoxicity in MDCK cells (CC_50_=1.64μM in Fig.1D) was less toxic to EBOV-mini-replicon supporting BSR-T7/5 cells (CC_50_=85.84μM). Thus, a significantly high SI=4746.11 was achieved by S416. We subsequently tested the inhibitory effects of DHODHi against Zika virus (**Fig. 2B**). EC_50_ values were 17.72, 0.268, 2.29 and 0.021μM for Teriflunomide, Brequinar, S312 and S416, respectively. Again, the selective index of S416 reached the top of SI=1087.62.

**Figure 2.**
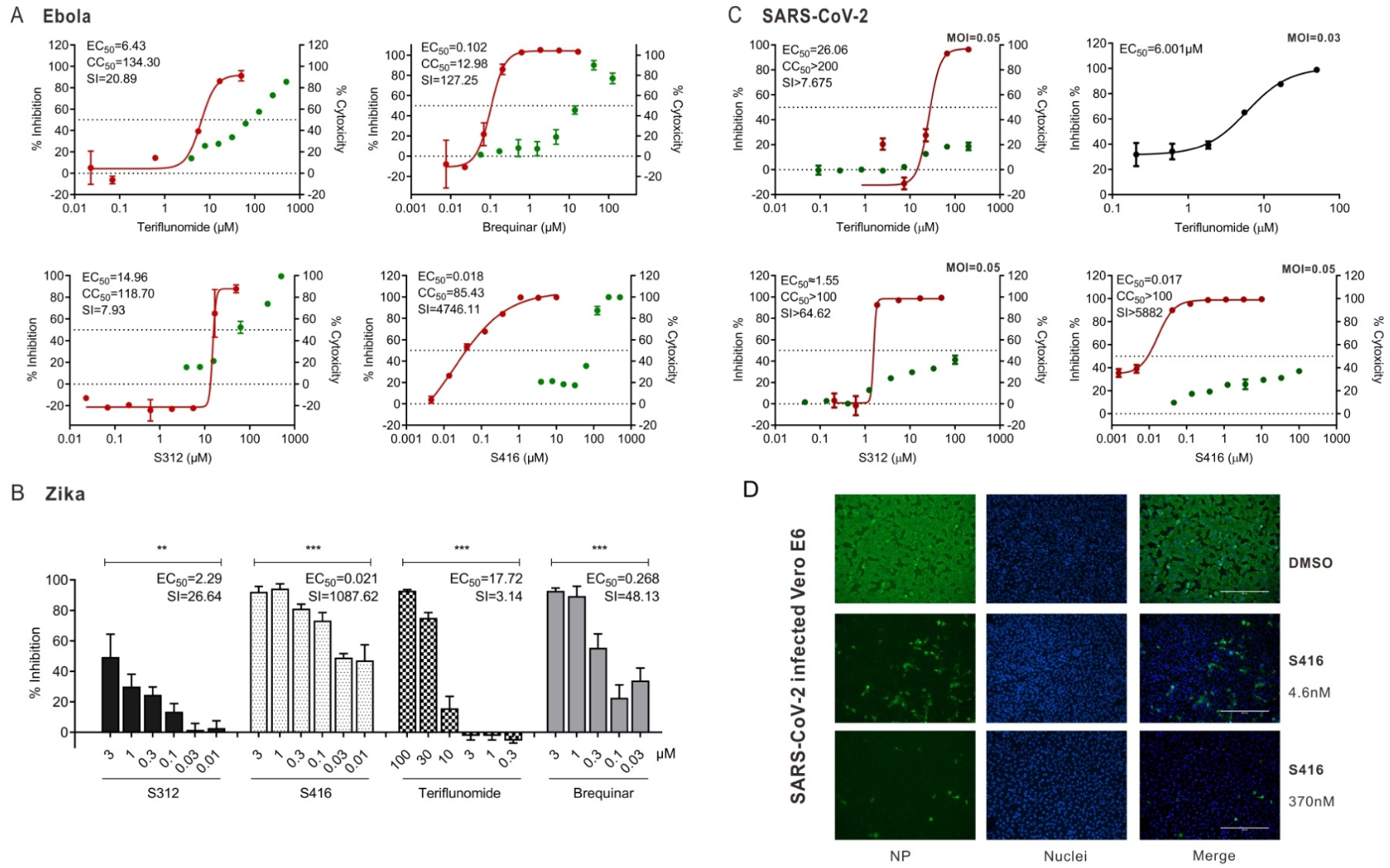
Broad-spectrum antiviral activities of DHODH inhibitors. **(A)** Anti-Ebola replication efficacy. BSR-T7/5 cells were transfected with the EBOV mini-genome replication system (NP, VP35, VP30, MG, and L) in the presence of increasing concentrations of Teriflunomide, Brequinar, S312 and S416 respectively. Inhibitory effects of these compounds (EC_50_) to EBOV mini-genome replication were determined using Bright-Glo Luciferase Assay (left-hand scale, red curve). CC_50_ of compounds were determined by analyzing BSR-T7/5 cell viability using CellTiterGlo Assay (right-hand scale, green curve). The results are presented as a mean of at least two replicates ± SD. **(B)** Anti-Zika virus efficacy. Huh7 cells were infected with Zika virus (MOI=0.05) for 4 hours and then treated with increasing concentrations of compounds Teriflunomide, Brequinar, S312 and S416 respectively. The viral yields in cell supernatants were then quantified by qRT-PCR to reflect the replication efficiency of Zika virus. **(C)** Anti-SARS-CoV-2 virus efficacy. Aliquots of Vero E6 cells were seeded in 96-well plates and then infected with Beta CoV/Wuhan/WIV04/2019 at MOI of 0.03. At the same time, different concentrations of the compounds were added for co-culture. Cell supernatants were harvested 48 h.p.i. and RNA was extracted and quantified by qRT-PCR to determine the numbers of viral RNA copies. **(D)** Immuno-fluorescence assay of SARS-CoV-2-infected cells. Vero E6 cells were infected with SARS-CoV-2 under the same procedure of C. Cells were fixed and permeabilized for staining with anti-viral NP antibody, followed by staining with Alexa 488-labeled secondary antibody. Green represents infected cells. Nuclei were stained by DAPI, and the merge of NP and nuclei were shown. Scale bar, 400uM. The results (B, C) are presented as a mean of at least three replicates ± SD. Statistical analysis, One-way ANOVA for (B). NS, *p* >0.05; *, *p* <0.05; **, *p* <0.01; ***, *p* <0.001.

When we prepared the manuscript, a severe outbreak of SARS-CoV-2 occurred in Wuhan in December 2019, we responded quickly to examine the antiviral activity of DHODHi against this new coronavirus. The data in **Fig. 2C** showed that all the DHODHi tested is low toxic to SARS-CoV-2 susceptible Vero E6 cells. Teriflunomide had a solid antiviral efficacy of EC_50_=26.06uM (at MOI=0.05, ∼2.4-fold stronger than Favipiravir [EC_50_=61.88μM]^29^) (**Fig. 2C upper right**), whereas its pro-drug Leflunomide showed less inhibition of EC_50_=63.56uM (data not shown). We therefore further did immuno-florescent assay to visualize the drug efficacy. To determine more carefully the efficacy of Teriflunomide, which can be transferred to clinical treatment of SARS-CoV-2 immediately as an approved drug, a bit low MOI of 0.03 (**Fig. 2C upper left**) were applied. In this condition, the EC_50_ of Teriflunomide could reach 6μM with SI>33, indicating that Teriflunomide with effective EC_50_ and SI values have all the potentials to treat SARS-CoV-2-induced COVID-19 disease as an ‘old drug in new use’ option.

Additionally, S312 and S416 exhibited ideal antiviral efficacies of EC_50_=1.55μM (SI>64.62) and EC_50_=0.017μM (extensively high SI>5882.4), respectively (**Fig. 2C lower panel**). Compared with our previous publication of Remdesivir (EC_50_=0.77μM, SI>129.87) and Chloroquine (EC_50_=1.13μM, SI>88.5)^29^, which are currently used in clinical trials against SARS-CoV-2, S416 had much greater EC_50_ and SI values (66.5-fold stronger than Chloroquine in EC_50_) against SARS-CoV-2. The data in **Fig. 2D** clearly showed that as little as 4.6nM (0.3EC_50_) of S416 can dramatically inhibit SARS-CoV-2 infections, while, increased drug concentration of 370nM (22EC_50_) could further eliminate viral infected cells. Thus, S416 turns to be the best efficient chemical so far against SARS-CoV-2 at the cellular level.

### S312 exhibited equivalent antiviral activities to Oseltamivir in influenza-A-virus-infected animals

In all the previous studies, inhibitors to DHODH or pyrimidine synthesis pathway were only active in vivo at low efficacy showing no obvious superior to DAA drugs. To test whether our potent DHODHi could refresh the important role of targeting DHODH in viral disease, we next explored *in vivo* efficacy of S312 by intranasally infecting BALB/c mice at a lethal dose of WSN (2LD_50_=4000pfu) or 2009 pandemic H1N1 (A/Sichuan/01/2009, SC09) (300pfu) virus. Treatment of S312 or Osel or combined treatment of ‘S312+Osel’ was given by the intraperitoneal (i.p.) route at 4h p.i. once per day from the beginning of infection to Day 14 to determine body weight changes and mortality (experimental procedure shown in **Fig. 3A**). The data in **Fig. 3B** showed that the bodyweights of mice from the DMSO-treated ‘virus group’ all dropped to less than 75% and died at D8 p.i.. DAA drug of Osel could indeed totally rescue all the mice from bodyweight loss and death. Equivalently, S312 (5mg/kg, red line) was also able to confer 100% protection and little bodyweight loss similar to Osel. Even S312 of 2.5mg/kg and 10mg/kg cold confer 75% protection and 50% protection, respectively. Considering the Cmax of S312 (≈15μM), we used both 5mg/kg and 10mg/kg in the following experiments. The results suggest that S312 of a modest dose (5mg/kg) would achieve the equivalent 100% protection to DAA drug when used from the beginning of infection.

**Figure 3.**
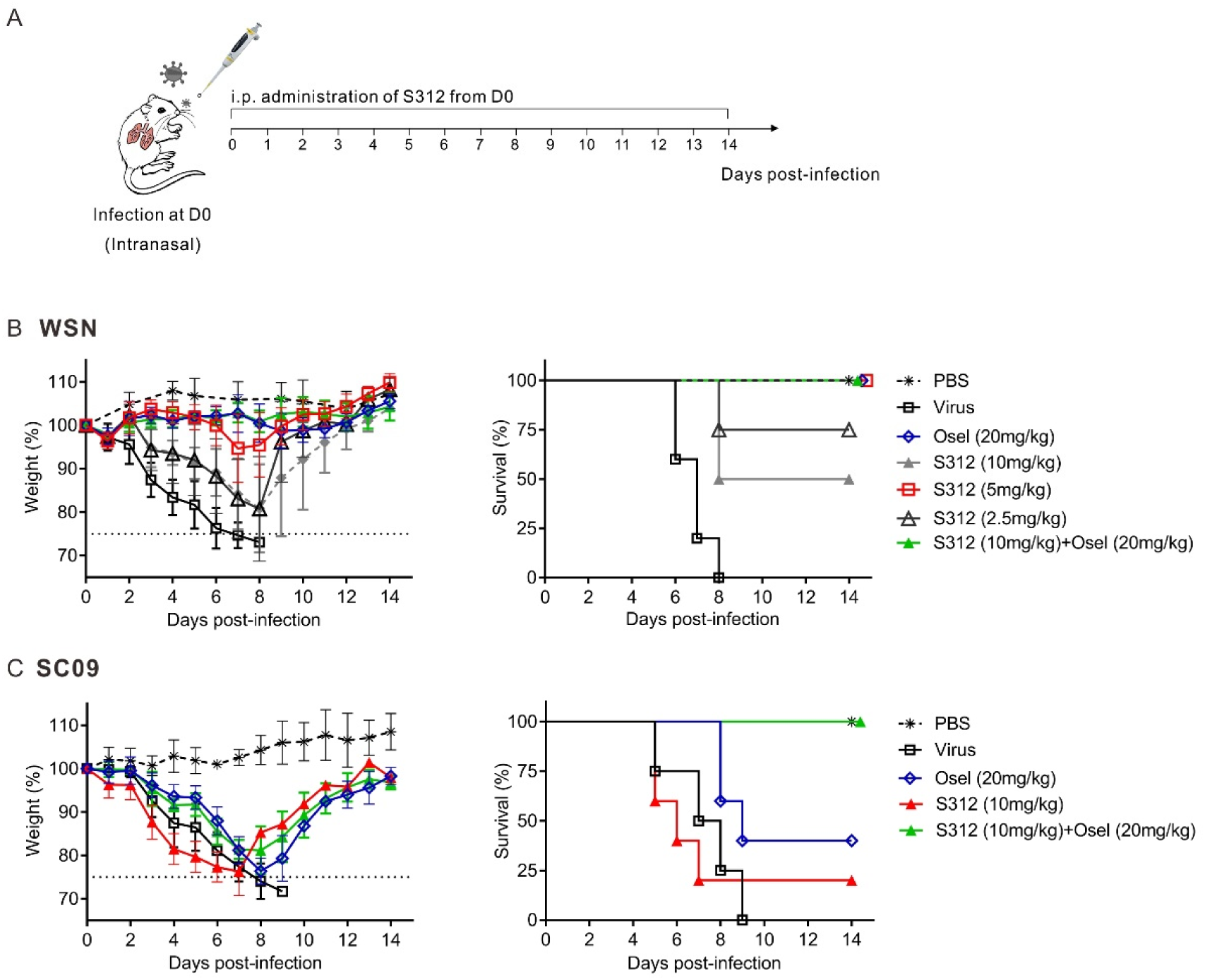
The *in vivo* antiviral activity of S312 in influenza-A-virus infected mice. **(A)** Diagram of the experimental procedure. **(B)** BALB/c mice were intranasal infected with 4000PFU of WSN virus and then intraperitoneal injected (i.p.) with PBS, S312 (2.5, 5, 10mg/kg), Oseltamivir (20mg/kg) and S312+Oseltamivir (10mg/kg+20mg/kg) once per day from D1-D14 respectively. The body weight and survival were monitored for 14 days or until body weight reduced to 75% (n = 4 mice per group). **(C)** Mice were inoculated intranasally with 600 PFU of A/SC/09 (H1N1) and then i.p. with S312 (10mg/kg), Oseltamivir (20mg/kg) and S312+Oseltamivir (10mg/kg+20mg/kg) once per day from D1 to D14. The body weight and survival were monitored until 14 days post-infection or when the bodyweight reduced to 75%. The dotted line indicates endpoint for mortality (75% of initial weight). The body weights are present as the mean percentage of the initial weight ±SD of 4-5 mice per group and survival curve were shown.

Except for broad active to different viruses, HTA drug such as DHODHi has another advantage over DAA drug to overcome drug-resistant. To prove this, we generated a current-circulating Oseltamivir-resistant NA^H275Y^ mutant virus (in WSN backbone) by reverse genetics (**Supplementary data Fig. 2A and 2B**). We found that the NA^H275Y^ virus did not respond to Osel (20mg/kg/day)-treatment at all but 2.5mg/kg/day of S312 can rescue 50% of mice from lethal infection of NA^H275Y^ virus (**Supplementary data Fig. 2C**). When mice were infected with a natural-isolated pandemic strain SC09, which is less sensitive to either Osel (40% protection) or S312 (20% protection) (**Fig. 3C**), we further observed 100% protection in combined treatment of S312+Osel, indicating that HTA and DAA drug combination can augment therapeutic effects.

The data in all refresh DHODH as an attractive host target in treating viral disease with equivalent efficacy to DAA drug, and is advantageous when facing DAA-drug-resistant viruses.

### DHODHi inhibit virus replication through interrupting the *de novo* pyrimidine synthesis pathway

To elucidate the essential role of DHODH in the viral replication cycle, we generated a DHODH^-/-^ A549 cell line by CRISPR-Cas9 gene knock-off (KO) technology (**Fig. 4A**). Unexpectedly, the cell proliferation rate was barely affected in DHODH^-/-^ cells indicating DHODH is not indispensable for cell growth at least for three days (72 hours) (**Supplementary Fig. 3**). By contrast, virus growth was largely inhibited in DHODH^-/-^ cells as compared to wild-type (WT) A549 cells with almost 1000-fold reduction of infectious particles at 72 hours post-infection (h.p.i.) (**Fig. 4B**). When S312 (5IC_50_) was added into the culture medium, dramatic reduction of virus growth only occurred in WT cells but not in DHODH^-/-^ cells (**Fig. 4C and 4D**). These results prove that virus growth but not the coincident cell growth requires DHODH activity, and antiviral action of S312 is implemented by targeting DHODH.

**Figure 4.**
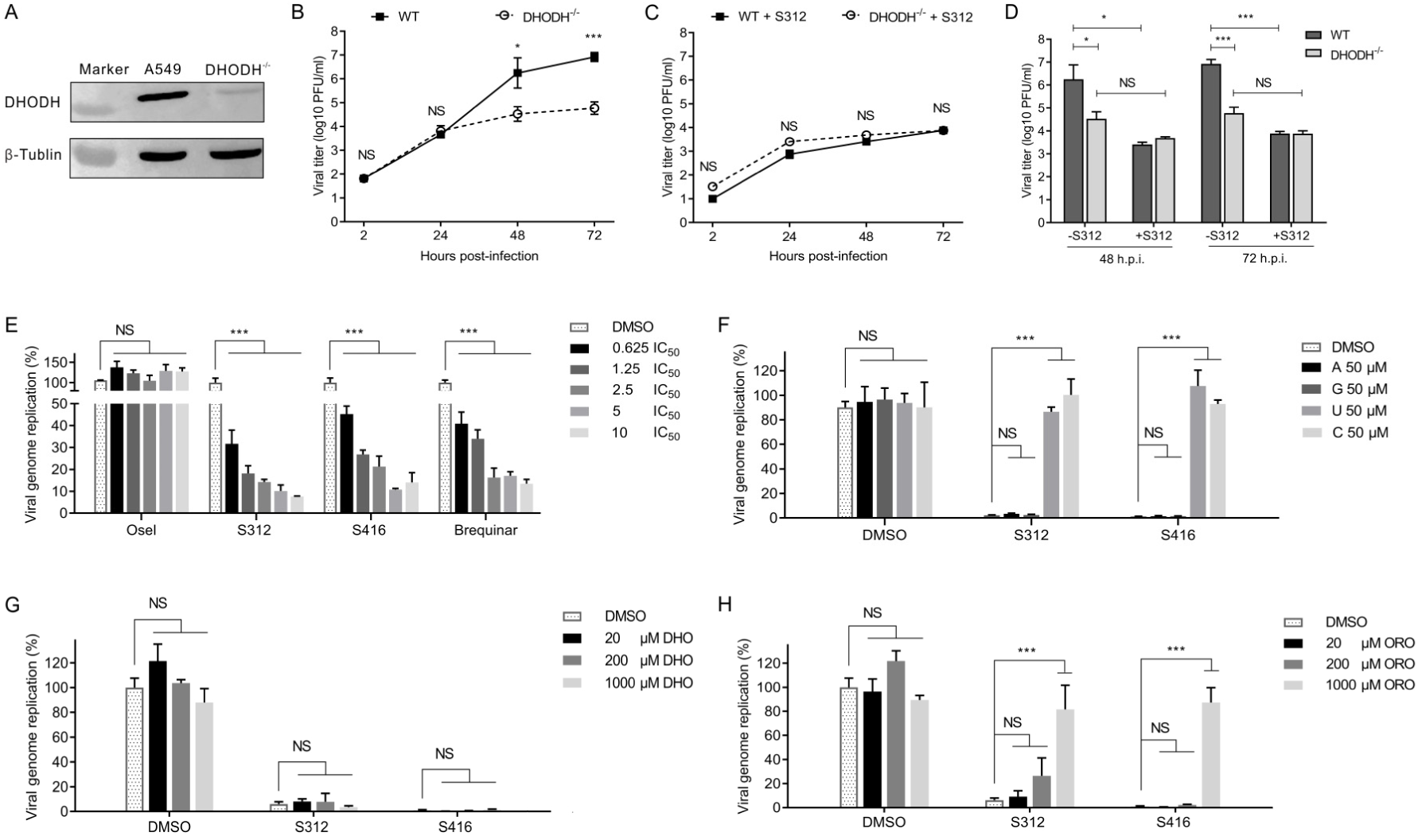
DHODHi inhibit virus replication through interrupting the *de novo* pyrimidine synthesis pathway. **(A)** The expression levels of DHODH protein in WT and DHODH^-/-^ A549 cells. **(B)** The growth curve of the WSN virus on A549 (WT or DHODH^-/-^) cells. A549 (WT or DHODH^-/-^) cells were infected with WSN virus with MOI of 0.01. The supernatants were assayed for viral titers at 2h, 24h, 48h, and 72h post-infection by plaque assay. **(C)** The same experimental procedures were performed as shown in (B) after adding S312 at a final concentration of 12.5 μM (5IC_50_). **(D)** Statistics of 48h and 72h of viral titers with or without S312 on A549 (WT or DHODH^-/-^) cells. **(E)** The 293T cells were co-transfected with the influenza virus minigenome plasmid system (PB1, PB2, PA, NP, pPoII-NP-luc, and pRLSV40). After 12 h.p.i., cells were treated with 2-fold serial dilutions of Oseltamivir, S312, S416, and Brequinar respectively. The luciferase activities were measured 24 h of post-treatment. **(F)** Effects of nucleotides addition on the antiviral efficacies of S312 and S416. The 10-fold IC_50_ of S312 (24 μM) or S416 (0.6 μM) and 50μM four nucleotides (Adenosine, Uridine, Cytidine, Guanosine) were added at the same time on 293T cells. After 24 h of treatment, the luciferase activities were measured. **(G and H)** Effects of addition of dihydroorotate (DHO) or Orotic acid (ORO) on the antiviral efficacies of S312 and S416. The luciferase activities were detected as above after treating with indicated concentrations of DHO or ORO. All results are presented as a mean of three replicates ± SD. Statistical analysis, two-way ANOVA for B, C, and D. One-way ANOVA for E, F, G and H. NS, *p* >0.05; *, *p* <0.05; **, *p* <0.01; ***, *p* <0.001.

The general virus growth cycle includes virus entry, viral genome replication, and virus release. To further validate virus genome replication is the major target of DHODHi, we used the influenza-A-virus mini-replicon system to quantify viral genome replication. Brequinar, another potent inhibitor of DHODH was included as a positive control^30^, whereas, Osel targeting influenza NA protein served as a negative control. The results in **Fig**.**4E** showed no inhibition on viral genome replication in the Osel-treated group, but there were obvious inhibitions on viral genome replication in both S312- and S416-treated groups as well as Brequinar-treated group in dose-dependent manners. Almost 90% of viral genome replication was suppressed by 10IC_50_ of S312 (24μM) and S416 (0.6μM). As DHODH catalyzes oxidation of dihydroorotate (DHO) to produce orotic acid (ORO) and finally forms UTP and CTP, we add four nucleotides (adenine nucleotide(A), guanine nucleotide(G), uracil nucleotide(U), and cytosine nucleotide(C)), DHO, ORO respectively to mini-replicon system to identify the target of S312 and S416. The results in **Fig. 4F** showed that the addition of 50μM either U or C could effectively rescue viral genome replication in S312-and S416-treated cells (as well as Brequinar-treated cells), whereas addition of neither A nor G changed the inhibitory effects. Moreover, supplement of DHODH substrate DHO cannot rescue viral genome replication (**Fig. 4G)**, but a supplement of DHODH product ORO can gradually reverse the inhibition effects of S312 and S416 (**Fig. 4H**). The results further confirm that compounds S312 and S416 inhibit viral genome replication via targeting DHODH and interrupting the fourth step in *de novo* pyrimidine synthesis.

### S312 is advantageous over the DAA drug to treat advanced and late-phase disease with decreasing cytokine/chemokine storm

It is documented elsewhere that DAA drugs such as Osel is only completely effective in the early phase of infection, optimally within 48 hours of symptom onset^31^. And till now, there is no approved drug to treat advanced influenza disease at the late phase specifically. We suppose that S312 could be effective in the middle or late phase of disease because it targets a host pro-viral factor of DHODH not affected by viral replication cycle. To test this, we compared the therapeutic windows of S312 and Osel in early (D3-D7), middle & late (D5-D9), severe late (D7-D11 or D6-D14) phases (workflow shown in **Fig. 5A**). When drugs were given in the early phase, both Osel-treatment and ‘Osel+S312’-combination-treatment conferred 100% protection (**Fig. 5B**). When drugs were given at the middle & late phase (**Fig. 5C**), single Osel-treatment wholly lost its antiviral effect with no surviving. However, S312-treatment could provide 50% protection, and drug combination reached to 100% protection. When drugs were given at severe late phase of disease that mice were starting dying (**Fig. 5D**), neither single treatment of Osel nor S312 can rescue the mice from death but combined treatment still conferred to 25% survival. To really show the advantage of S312 in treating severe disease, we additionally treated the mice a bit early before dying around 80% of initial weights (D6-D14) with a more optimal dose of S312 (5mg/kg). The data in **Fig. 5E** showed that S312 rescued 50% of mice from severe body-weight losses, and combined treatment coffered additional 25% survival. These results once again highlight that S312 has remarkable advantages over Osel to treat severe diseases at the late phase, and its therapeutic effectiveness could even be improved when S312 was combined with DAA drug.

**Figure 5.**
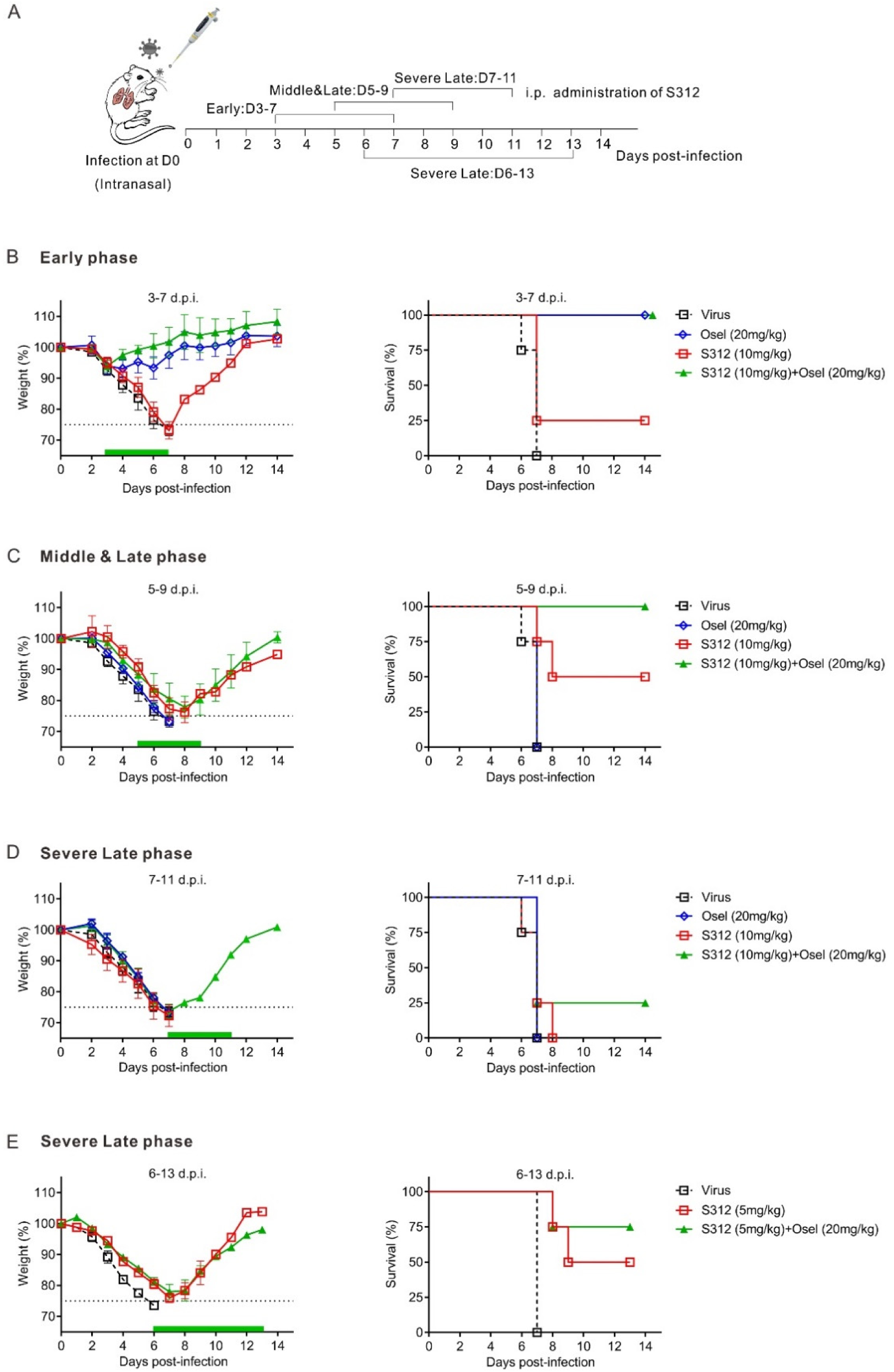
S312 is more effective at the late and severe infection phase as compared to DAA drug Oseltamivir. **(A)** Diagram of the experimental procedure. **(B-E)** BALB/c mice were inoculated intranasally with 4000PFU of WSN virus and then i.p. with S312 (10mg/kg), Oseltamivir (20mg/kg), or S312+Oseltamivir (10mg/kg+20mg/kg) once per day from D3-7 **(B)**, D5-9 **(C)**, D7-11 **(D)**. Another groups of S312 (5mg/kg) or S312+Oseltamivir (5mg/kg+20mg/kg) were given i.p. once per day from D6 to D13 in **E**. The green bars indicate the period of drug administration. The body weight and survival were monitored until 14 days post-infection or when the bodyweight reduced to 75%, respectively (n =4-5 mice per group). The dotted line indicates endpoint for mortality (75% of initial weight). The body weights are present as the mean percentage of the initial weight ±SD of 4-5 mice per group and survival curve were shown.

It is known that severe acute infections including influenza and COVID-19 always induce pathogenic immunity as cytokine/chemokine storms. Leflunomide and Teriflunomide are already clinically used in autoimmune disease to inhibit pathogenic cytokines and chemokines. We, therefore, suspect that DHODHi should also be anti-cytokine-storm in viral infectious disease. BALF from either Osel- or ‘S312+Osel’-treated mice were collected at D14 in an independently repeated experiment with a lower infection dose. A parallel body weights excluded differences in virus load (**Supplementary Fig. 4A**). The data in **Supplementary Fig. 4B** showed that the pathogenic inflammatory cytokines in ‘S312+Osel’-treated group was largely reduced as compared to Osel-treated mouse in the levels of IL6, MCP-1, IL5, KC/GRO(CXCL1), IL2, IFN-γ, IP-10, IL9, TNF-α, GM-CSF, EPO, IL12p70, MIP3α and IL17A/F (listed in the order of reduce significance).

The results in all provide striking information that DHODH inhibitors are effective in infected animals not only by inhibiting virus replication (Shown in Fig. 3 and Fig. 4) but also by eliminating excessive cytokine/chemokine storm. Usage of DHODHi could finally benefit to advanced disease in late infection.

## Discussion

Many different RNA viruses had caused epidemics and pandemics in the past 20 years, from avian influenza virus (AIV) H5N1 in 1997, SARS-CoV in 2002, pandemic H1N1 in 2009, MERS-CoV in 2012, AIV H7N9 in 2013, Ebola virus in 2013, Zika virus in 2016, to SARS-CoV-2 in 2019. Many of them are still prevalent in certain countries and SARS-CoV-2 continues causing huge damage to China and worldwide. The conventional one-bug-one-drug paradigm is insufficient to deal with the challenges of these emerging and re-emerging pathogenic viruses^32^, moreover, it is too late to develop drugs against each virus behind epidemics^33^. Thus, it is imperative to develop new ideas, highly effective, broad-spectrum antiviral agents that can treat various viral infections. In this study, we applied DHODH inhibitors including a computer-aided designed compound S312 into viral infectious disease. We found that direct-targeting DHODHi are broad-spectrum antiviral both in cell culture and *in vivo*. The candidate S312 had further advantage to be used in infected animals with low toxicity and high efficiency. Moreover, S312 can rescue severe influenza infection by limiting inflammatory cytokine storm *in vivo*.

DHODH is a rate-limiting enzyme catalyzing the fourth step in pyrimidine *de novo* synthesis. It catalyzes the dehydrogenation of dihydroorotate (DHO) to orotic acid (ORO) to finally generate Uridine (U) and Cytosine (C) to supply nucleotide resources in a cell. Under normal conditions, nucleotides are supplied via both *de novo* biosynthesis and salvage pathways, the latter of which is a way of recycling pre-existing nucleotides from food or other nutrition. However, in virus-infected cells, a large intracellular nucleotide pool is demanded by rapid viral replication. It is therefore reasonable that *de novo* nucleotides biosynthesis rather than salvage pathway is more critical for virus replication. Our data indeed show that virus replication is largely restricted when the DHODH gene was knocked off even with a complete culture medium. By contrast, cell growth was not affected by lacking DHODH at all, indicating that *de novo* nucleotides biosynthesis is not indispensable in normal cell growth without infection at least for days. **More interestingly**, we notice that compared with DNA viruses, RNA viruses need unique UMP but not TMP in their genomes. UMP is the particular nucleoside produced by DHODH, which means RNA viruses might be more sensitive to DHODH activity. SARS-CoV-2, for instance, has around 32% of UMP in its genome explaining why DHODHi are effective and superior to SARS-CoV-2. Nevertheless, the comparison between different viruses is worth to be studied in the future.

Although several DHODHi have been documented to be antiviral by high-throughput screening^34-37^. Most of these compounds are still at cell culture level with unknown *in vivo* efficacy. Therefore, the development of broad-spectrum antiviral agents targeting DHODH is still an exciting avenue in antiviral research. S312 and S416 present more potent inhibition and favorable pharmacokinetic profiles, moreover, the half-lives of S312 and S416 (8.20 and 9.12 h, respectively) are much shorter and more appropriate than that of Teriflunomide, indicating that they may have less possibility to bring toxic side effects from drug accumulation in the body. Strikingly, S312 showed active effects *in vivo* in lethal dose infection of influenza A viruses not only when used from the beginning of infection but also in the late phase when DAA drug is not responding anymore. Another surprise is the high SI value of S416 to against Zika (SI=1087.62), Ebola (SI=4746.11), and the current SARS-CoV-2 (SI>5882). These data interpreted that S416 is highly promising to develop further as it should be to S312. The extremely high SI of **S416** may be due to its high binding affinity and favorable occupation of the ubiquinone-binding site of DHODH with faster-associating characteristics (*k*_on_ = 1.76×10^6^ M^-1^s^-1^) and slower dissociating binding characteristic (*k*_off_=2.97×10^−3^ s^-1^), which will reduce the possibility of off-target *in vivo*.

Acute viral infections usually cause severe complications associated with hyper induction of pro-inflammatory cytokines, which is also known as “cytokine storm” firstly named in severe influenza disease^38,39^. Several studies showed that lethal SARS patients expressed high serum levels of pro-inflammatory cytokines (IFN-γ, IL-1, IL-6, IL-12, and TGFβ) and chemokines (CCL2, CXCL10, CXCL9, and IL-8) compared to uncomplicated SARS patients^40-43^. Similarly, in severe COVID-19 cases, ICU patients had higher plasma levels of IL-2, IL-7, IL-10, GSCF, MCP1, MIP1A, and TNFα compared to non-ICU patients^44^. Moreover, A clinical study of 123 patients with COVID-19 showed that the percentage of patients with IL-6 above normal is higher in severe group^45^. In terms of treatment, immunomodulatory agents can reduce mortality and organ injury of severe influenza. However, these immunomodulatory are mostly non-specific to viral infection but rather a systemic regulation, such as corticosteroid, intravenous immunoglobulin (IVIG) or angiotensin receptor blockers^46-49^. Leflunomide and its active metabolite Teriflunomide have been approved for clinical treatment for excessive inflammatory diseases such as rheumatoid arthritis and multiple sclerosis^50^. Our data once again proved that DHODHi could further reduce cytokine storm than DAA drugs when using influenza-A-virus infected animal as a model. We believe that a similar immune-regulating role of DHODHi will exist in COVID-19 patients. Thus, by targeting DHODH, the **single key enzyme** in viral genome replication and immune-regulation, **a dual-action** of DHODH can be realized in fighting against a broad spectrum of viruses and the corresponding pathogenic-inflammation in severe infections. We hope our study may quickly and finally benefit the patients now suffering from severe COVID-19 and other infectious diseases caused by emerging and re-emerging viruses.

## Materials & Methods

### Cell lines, virus, and drugs

MDCK, A549, Vero E6, Huh7 cells were obtained from the American Type Culture Collection (ATCC). 293FT cells were kindly provided by Paul Zhou (Institute Pasteur of Shanghai, China). BSR T7/5 cells stably expressing the T7 RNA polymerase gene were kindly provided by Gang Zou (Institute Pasteur of Shanghai, China). MDCK, A549, Vero E6, Huh7, 293FT cells were cultured in DMEM (Gibco) supplemented with 10%FBS (Gibco) and 1% P/S (Gibco). BSR T7/5 cells were cultured in GMEM (Gibco) supplemented with 10%FBS (Gibco), 1% L-Glutamine (Gibco), 2% MAA and 1% P/S (Gibco). The A/WSN/33 virus was generated by reverse genetics as previously described^51^. The A/Sichuan/1/2009(H1N1)(SC09), A/Jiangxi-Donghu/312/2006(H3N2), A/Guangzhou/333/99(H9N2) were kindly provided by Prof. Yuelong Shu (Chinese National Influenza Center). Zika virus SZ-WIV001 (KU963796) was kindly provided by Bo Zhang from Wuhan Virology Institute of CAS^52^. DHODH inhibitors S312 and S416 were synthesized using our previously reported synthetic routes^27,53^. Leflunomide, Teriflunomide, Brequinar, Oseltamivir, Adenosine, Uridine, Cytidine, Guanosine, Orotic acid, Dihydroorotate were purchased from Sigma-Aldrich. DHODH knock-out engineered A549 cell line was generated by CRISPR/Cas9 gene-editing system. The following four group primers for sgRNA were synthesized by Generay Company (Shanghai, China), and cloned into lentiCRISPRv2 vector provided by Dimitri Lavillette (Institute Pasteur of Shanghai, China). 293FT cells were then co-transfected with lentivirus packaging plasmids (lenti-CRISPR v2-DHODH-sgRNA, psPAX2, and PMD2.G) by Lipofectamine® 2000 (ThermoFisher) for 48 hours. The lentivirus supernatants were harvested after centrifugation and filtration to (then mix supernatants of four sgRNAs) to transduce 1.5 × 10^6^ A549 cells into a 6-well plate for 72 h at 37 °C. Cells were expanded into 10 mm plates and grown in the presence of puromycin (5 μg/mL) for 3–5 days to select sgRNA-positive cells. Thereafter, single-cell sorting was performed in a 96-well plate to grow single clones, and western blot using the anti-DHODH antibody (Santa Cruz) was performed to detect the expression level of DHODH. The best KO cells with the lowest residue DHODH level were finally used. The primers of DHODH sgRNAs were listed below:

DHODH_sg1 (F: 5’-CACCGACACCTGAAAAAGCGGGCCC-3’, R: 3’-CTGTGGACTTTTTCGCCCGGGCAAA-5’)

DHODH_sg2 (F: 5’-CACCGCGTGGACGGACTTTATAAGA-3’, R: 3’-CGCACCTGCCTGAAATATTCTCAAA-5’)

DHODH_sg3 (F: 5’-CACCGCGCAGAAGGGGTGCGCGTAC-3’, R: 3’-CGCGTCTTCCCCACGCGCATGCAAA-5’)

DHODH_sg4 (F: 5’-CACCGTCAAAGAGTTGGGCATCGAT-3’, R: 3’-CAGTTTCTCAACCCGTAGCTACAAA-5’)

### Isothermal titration calorimetry (ITC)

ITC measurements were carried out at 25 °C on a MicroCal iTC200 (GE Healthcare). For the titration of an inhibitor to DHODH, both were diluted using the buffer (20 mM HEPES, pH8.0, 150 mM KCl, 10% Glycerol and 10% DMSO). The concentration of DHODH in the cell was 20 µM, and the concentration of inhibitor in the syringe was 150 µM for S312 or 100 µM for S416. All titration experiments were performed by adding the inhibitor in steps of 2 µL. The data were analyzed using Microcal origin software by fitting to a one-site binding model.

### Surface plasmon resonance (SPR)

Surface plasmon resonance experiments were performed with a BIAcore T200 (GE Healthcare) according to our previous work ^53^. The running buffer contained 1.05×PBS, 0.005% (vol/vol) surfactant P20, pH 7.4, and 1% DMSO. The purified DHODH, which was diluted in sodium acetate solution (pH 5.5) with a final concentration of 30 μg/mL, was immobilized on a CM5 sensor chip by amine coupling. All analyte measurements were performed at a flow rate of 30 μL/min. The analyte was diluted in the running buffer from the top concentration. Data processing and analysis were performed using BIAevaluation 1.1 software.

### Anti-influenza virus activity

Based on CellTiter-Glo® Cell Viability Assay (Promega), anti-influenza virus activities of compounds were evaluated. For compound toxicity assays (CC_50_), MDCK in a 96-well plate at a density of 2×10^4^cell/well were cultured in the present of serial dilutions of compounds in infection solution (DMEM + 0.2% BSA + 25 mM HEPES + 1 μg/mL TPCK). After 72 hours of incubation in a 37 °C, 5% CO_2_ incubator, 25 µl of CellTiter-Glo® reagent was added to each well, and then record the fluorescence (Luminescence) value to reflect cell viability. For compound effective assays (EC_50_), the same numbers of cells were seeded in a 96-well plate and were infected by 50 µL of a 20 TCID_50_ virus suspension and another 50 µL of drug dilutions. The cells were then incubated at 37 °C until the virus control group reached a 75%-100% cytopathic effect. Cell viability was quantified as above. All drug concentrations were performed at least three replicates. Data were processed with Graphpad prism software to calculate EC_50_ and CC_50_ values of the compounds.

### Anti-Zika virus activity

Aliquots of Huh7 (1×10^4^ cells/well) were seeded into 12-well plates and infected with the Zika virus (MOI=0.05). At 4h post-infection, the medium was removed, and cells were treated with appropriate concentration of compounds in infection medium (DMEM+2% FBS+25 mM HEPES). After 7 days, the supernatant was collected and RNA was extracted by TRIzol Reagent (Invitrogen) and submitted to reverse transcription-quantitative PCR with the PrimeScript RT reagent kit (TaKaRa). The primers to detect Zika-specific RNA were listed below: Zika virus NS3 (F: 5’-CTCCAGGATGCAAGTCTAAG -3’, R: 5’-ACCCAGCAGGAACTTCAGGA -3’). All drug concentrations were performed at least three replicates. Data were processed with Graphpad prism software to calculate EC_50_ and CC_50_ values of the compounds.

### Bright-Glo Pharmacodynamic assay for Ebola Replicon

Ebolavirus must be operated in the BSL-4 laboratory. To reduce the biological safety risks, the Ebolavirus replicon system was chosen for antiviral efficacy assay^54^. BSR T7/5 cells (T7 RNA polymerase stable expressing cell) were seeded in 96-well plates and co-transfected with p3E5E-Luc (25 ng), pCEZ-NP (25 ng), pCEZ-VP35 (25 ng), pCEZ-VP30(15 ng), and pCEZ-L (150 ng) per well. Afterward, 50 μL of drug dilutions were added into each well and cultured in at 37 °C, 5% CO_2_ for 24 hours. Replicon efficacy was determined by adding 50 μL/well of Bright-Glo reagent to record the fluorescence (Luminescence) values. All drug concentrations were performed at least three two replicates. Data were processed with Graphpad prism software to calculate EC_50_ and CC_50_ values of the compounds.

### Detection of anti-SARS-CoV-2 efficacy by real-time quantitative PCR

Vero E6 cells seeded on 48-well plates were infected with virus 2019BetaCoV/Wuhan/WIV04/2019 isolated by Wuhan Institute of Virology, Chinese Academy of Sciences at MOI of 0.03. At the same time, different concentrations of the drugs were added for co-culture, and DMSO dilution was used as a negative control. Cell supernatants were harvest 24 hours post-infection and RNA was extracted by a viral RNA extraction kit (Takara, Cat no. 9766). The number of viral copies was detected by real-time quantitative reverse transcription PCR (qRT-PCR) (Takara Cat no.RR820A) to reflect the replication efficiency of the virus. Data were processed by Graphpad prism software to calculate EC50 of the compound. The cytotoxicity of the tested drugs on cells was determined by CCK8 assays (Beyotime, China).

### IFA assay of SARS-CoV-2

To detect viral protein expression in Vero E6 cells, cells were fixed with 4% paraformaldehyde and permeabilized with 0.5% Triton X-100. The cells were then incubated with the primary antibody (a polyclonal antibody against the NP of a bat SARS-related CoV) after blocking, followed by incubation with the secondary antibody (Alexa 488-labeled goat anti-rabbit, Abcam).

### Virus growth curve and plaque assay

To determine the viral growth curve, A549 (WT and DHODH^-/-^) cells were infected with the indicated viruses at an MOI of 0.01. The supernatant of infected cells was collected at different time points post-infection and titrated by plaque assay on MDCK cells. The viral plaque assay with immunostaining was performed as previously described^55^. Briefly, MDCK cells cultivated in 24-well plates were infected with WSN virus. After 0.5 hours of incubation at 37 °C in 5% CO_2_, the virus medium was removed, and 1 ml of 2.4% Avicel (FMC Corporation) medium in the presence of indication drugs was added. Cells were incubated for another 48 hours at 37 °C in 5% CO_2_ and then fixed with 4% PFA. Immunostaining was performed using an anti-NP polyclonal antibody conjugated with HRP (Antibody Research Center, Shanghai Institute of Biological Science). True Blue substrate (KPL) was added to visualize the plaques.

### Influenza mini-replicon system and DHODH substrates assay

The 293T cells were seeded into 24-well plates at 1×10^5^ cells per well and were transfected with influenza A/WSN/33 virus PB1-, PB2-, PA- and NP-expressing plasmids (100 ng each), the influenza virus-specific RNA polymerase I driven firefly luciferase reporter (pPoII-NP-luc) (100 ng, provide by Han-D Klenk, Marburg University), and the Renilla luciferase reporter pRLSV40 10 ng, Promega). After 12 hours, the transfected 292T cells were treated with different concentrations of the test compounds or exogenous nucleosides (Adenosine, Uridine, Cytidine, Guanosine) or orotic acid or dihydroorotate. Cells were harvested after another 24 hours and firefly luciferase and Renilla luciferase expression were determined using the Dual-Luciferase Assay Kit (Promega).

### Mice infection and drug treatment

BALB/c female mice, 6-8 weeks old, were purchased from LingChang Company and bred in the specific pathogen-free animal facility. All animal experiments were approved by the Institutional Animal Care and Use Committee of Institut Pasteur of Shanghai and Wuhan University. For animal infection studies, mice were anesthetized using diethyl ether and intranasally challenged with virus suspension, i.e. 4000 PFU of A/WSN/33 H1N1 in 50ul of PBS and 600 PFU of A/SC/09 (H1N1) in 30ul of PBS, respectively. Diluted compounds were given by intraperitoneal (i.p) injection once a day. The drug treatment was initiated on days 0, 3, 5, 6, 7 post-infection respectively and continued for several days. Animal weight and survival were monitored daily, and mice were euthanized until the end of the experiment or when body-weight lost more than 25%.

## Data availability

The protein structure data has been uploaded to the Protein Data Bank with accession number 6M2B.

## Acknowledgments

This work was supported in part by the National Key Research and Development Program (grant 2018FYA0900801, 2018ZX10101004003001, 2016YFA0502304), the National Natural Science Foundation (NSFC) of China (grants 31922004, 81825020, 81772202), the National Science & Technology Major Project “Key New Drug Creation and Manufacturing Program” China (grant 2018ZX09711002), Application & Frontier Research Program of Wuhan Government (2019020701011463). Honglin Li is also sponsored by the National Program for Special Supports of Eminent Professionals and National Program for Support of Top-Notch Young Professionals. We thank Dr. Zhengli Shi for reviewing our manuscript. We also thank our group members of SARS-CoV-2 working group in State Key Laboratory of Virology, Wuhan University, who work tightly together during this new virus outbreak inside and outside of Wuhan city for their research spirits and courage, especially to Fang Liu, Qi Zhang, Ming Guo, Yuan Liu, Yan Zhang, Ying Zhu for their efforts. We are grateful to Taikang Insurance Group Co., Ltd and Beijing Taikang Yicai Foundation for their great supports to this work.

## Author Contributions

K.X. and H.L. conceived the project and designed the experiments; K.X., R.X., LK.Z., S.L., M.D., Y.W., Y.Z., Y.W., W.S., X.J., J.S., Y.T., L.X., Z.S., LL.Z. performed the experiments; D.L., K.L., G.X, C.Y., Y.L., Z.Z., R.W., G.Z. provide materials or technological supports, K.X., H.L., R.X., LK.Z., S.L., Y.W., Y.Z., Y.T., L.X., Z.S., K.L., G.X, Z.Z., R.W. and LL.Z., analyzed and discussed the data; K.X., H.L., R.X., LK.Z., S.L., Y.W., Y.Z., K.L., and LL.Z. wrote the manuscript.

