## Supplementary Tables and figures for "Novel and potent inhibitors targeting DHODH, a rate-limiting enzyme in *de novo* pyrimidine biosynthesis, are broad-spectrum antiviral against RNA viruses including newly emerged coronavirus SARS-CoV-2"

### Supplementary Materials and Methods

#### Crystallization and structure determination

The purified DHODH was concentrated to 20 mg/mL and then was incubated with 1 mM S416 and 2 mM DHO for 2h. Co-crystallization of DHODH with S416 was performed at 20 ºC using the vapor diffusion method as previously described [14,41]. X-ray diffraction data were collected at 100 K on beamline BL17U1 at the Shanghai Synchrotron Radiation Facility (SSRF). The raw data were processed using the MOSFLM and SCALA programs from the CCP4 suite [42,43]. The complex structure of DHODH-S416 was determined by molecular replacement using PDB entry 4LS1 (without ligands and water molecules) as the search model. Structure refinement was conducted with REFMAC5 [44]. Coot was used for the interpretation of electron density maps and model building [45]. Data collection and refinement statistics were listed in Supplementary Table 1.

#### Drug *in vivo* toxicity study

Mice were randomly assigned to five groups with one vehicle group and the other four groups orally administered S312 of 50, 100, 500 and 1000 mg/kg (n=4), respectively. Single-dose of S312 was administered on the first day of the experiment. During the experiment, all animals were clinically observed every day for 15 days for toxic signs. Bodyweight and food intake were recorded every two days. At the end of the experiment, all surviving animals were perfused, and samples of heart, liver, lung, and kidney were collected. Then the tissues were fixed in 4% paraformaldehyde, dehydrated with sucrose solution, and embedded in optimum cutting temperature. Frozen tissues sections (10 mm) were examined under a Nikon optical microscope at 100x magnification with standard hematoxylin and eosin staining.

#### Cytokine and chemokine measurements

BALB/c female mice, 6-8 weeks old, were anesthetized and intranasally challenged with 4000 PFU of A/WSN/33 H1N1 in 50 µL of PBS at day 0. Afterward, mice were weighed daily and administrated daily by intraperitoneal injection until the bodyweight dropped to about 80%. When the weight of the mice had dropped to 75% or the experiment finished, mice were euthanized and the bronchoalveolar lavage fluid was taken as previously published[31]. The samples were frozen at −80°C until subsequent analysis. The samples of lavage fluid were centrifuged at 3000 g for 10 min at 4 °C and the cytokines and chemokines were measured using the U-PLEX Biomarker Group 1 (ms) assays. All assays were performed according to manufacturer's instructions, tests were performed in duplicates, and without alterations to the recommended standard curve dilutions[32,33].

**Supplementary Table 1**.

Data collection and refinement statistics for the DHODH-S416 co-crystal structure

| **Data collection** | |
|---|---|
| Wavelength (Å) | 0.97852 |
| Space group | $P3_221$ |
| Cell dimensions | |
| *a*, *b*, *c* (Å) | 91.04, 91.04, 122.18 |
| $\alpha$, $\beta$, $\gamma$ (°) | 90.00, 90.00, 120.00 |
| Resolution (Å) | 48.34-1.76 (1.86-1.76) |
| $R_{merge}$ (%) | 10.7 (44.2) |
| $I / \sigma(I)$ | 15.0 (5.4) |
| Completeness (%) | 100 (100) |
| Redundancy | 10.3 (9.9) |
| **Refinement** | |
| Resolution (Å) | 48.34-1.76 (1.86-1.76) |
| No. of reflections used in refinement | 55638 |
| $R_{work}$ / $R_{free(\%)}$ | 16.7/19.3 |
| Total atoms | 3059 |
| Protein atoms | 2847 |
| Ligand (FMN/ORO/S416) atoms | 67 |
| Water | 145 |
| Wilson B-factor (Å$^2$) | 16.6 |
| Average B, all atoms (Å$^2$) | 20.0 |
| R.m.s. deviations | |
| Bond lengths (Å) | 0.010 |
| Bond angles (°) | 1.742 |
| Ramachandran plot | |
| Preferred regions (%) | 96.31 |
| Allowed regions (%) | 2.84 |
| Outliers (%) | 0.85 |
| **PDB code** | 6M2B |

*Values in parentheses are for the highest-resolution shell.

**Supplementary Table 2.**

Bioactivity and binding properties of DHODH inhibitors.

| Compd. | $\Delta H$ (kJ/mol) | $-T\Delta S$ (kJ/mol) | $\Delta G$ (kJ/mol) | $k_{on}$ ($M^{-1}s^{-1}$) | $k_{off}$ ($s^{-1}$) | $K_D$ (M) |
|---|---|---|---|---|---|---|
| **S312** | -38.83 | -6.23 | -45.06 | $3.12\times10^{5}$ | $6.33\times10^{-3}$ | $2.03\times10^{-8}$ |
| **S416** | -33.51 | -13.22 | -46.74 | $1.76\times10^{6}$ | $2.97\times10^{-3}$ | $1.69\times10^{-9}$ |

**Supplementary Table 3.**

Pharmacokinetics profiles of DHODH inhibitors S312 and S416.

| **Parameter**[a] | **S312**[2] | **S416** |
|---|---|---|
| IV dose (mg/kg) | 1 | 1 |
| $C_0$ (μg/L) | 34478.30±18771.00 | 6709.82±188.11 |
| $AUC_{0-\infty}$ (μg*h/L) | 23318.81±4501.24 | 47558.65±6150.08 |
| Cl (L/h/kg) | 0.04±0.01 | 0.02±0.00 |
| $Vd_{ss}$ (L/kg) | 0.35±0.02 | 0.14±0.02 |
| PO dose (mg/kg) | 10 | 10 |
| $T_{1/2}$ (h) | 8.20±1.72 | 9.12±2.91 |
| $AUC_{0-\infty}$ (μg*h/L) | 53047.73±11030.40 | 362768.43±132144.96 |
| $C_{max}$ (μg/L) | 5310.50±1910.47 | 24379.63±7533.42 |
| $T_{max}$ (h) | 1.00±0.87 | 4.00±0.00 |
| *F* (%) | 22.75±4.73 | 76.28±27.79 |

[a] Compounds were dosed to an equal number of male Sprague-Dawley rats in IV and PO administration respectively ($n$ = 3).

**Supplementary Fig. 1**

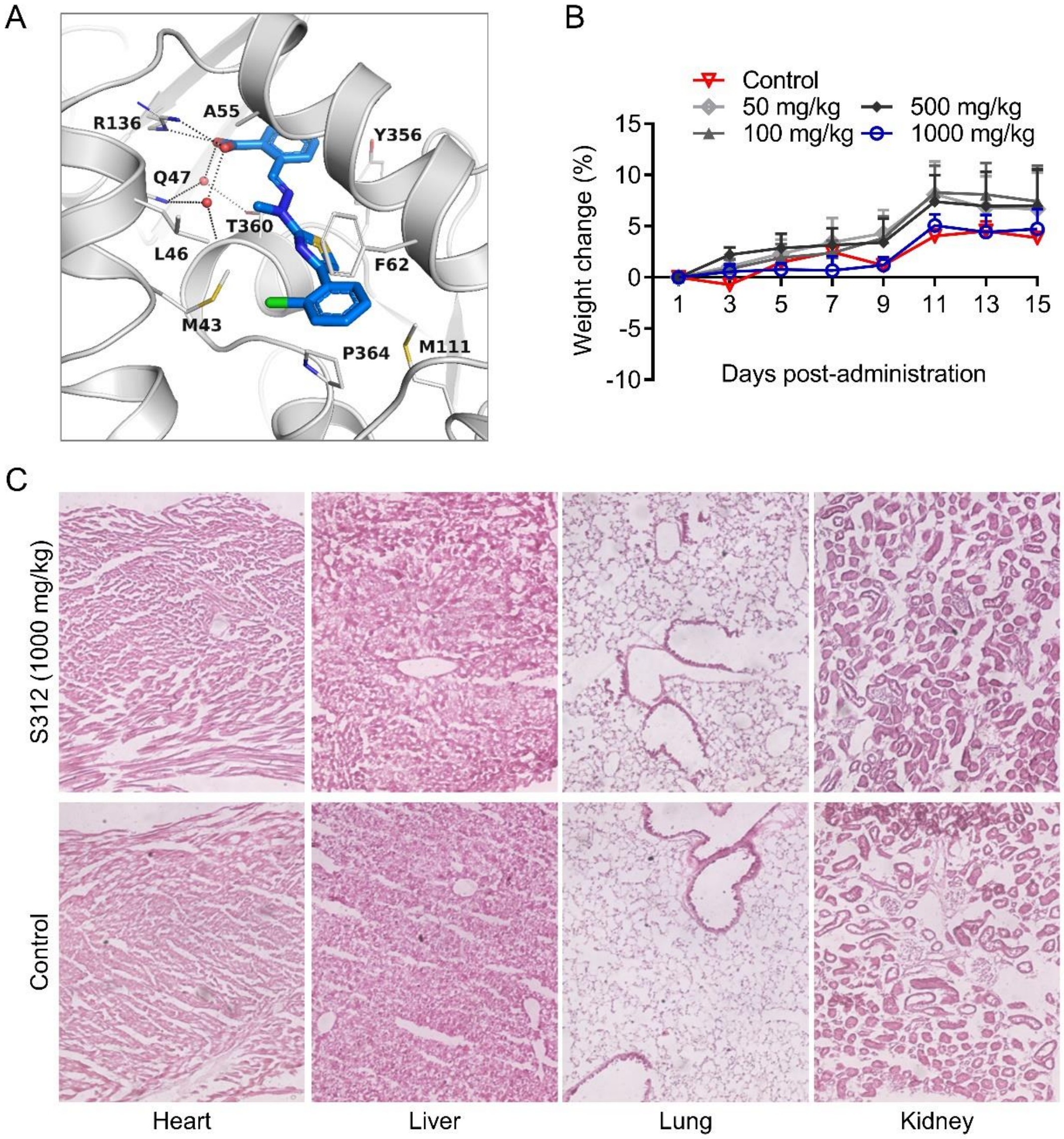


**Supplementary Data Fig. 1. Binding mode of S416, and the acute toxicity study of S312. (A)** The binding mode of S416 to DHODH in the X-ray complex crystal structure (PDB ID: 6M2B). Key residues are shown as gray thin sticks. Hydrogen bonds are displayed as black dashed lines. Water molecules are depicted as small red balls. **(B)** Bodyweight changes of the mice orally treated with S312 (50, 100, 500 and 1000 mg/kg) in the acute toxicity study. No significant differences in body weight changes were observed. The results were presented as means ± SEM. Statistical analysis: Student's t-test. **(C)** Hematoxylin and eosin stains of various tissues under an optical microscope for histopathological morphology analysis at 15 days after oral administration of 1000 mg/kg S312 in ICR mice. No abnormalities were observed.

**Supplementary Fig. 2**

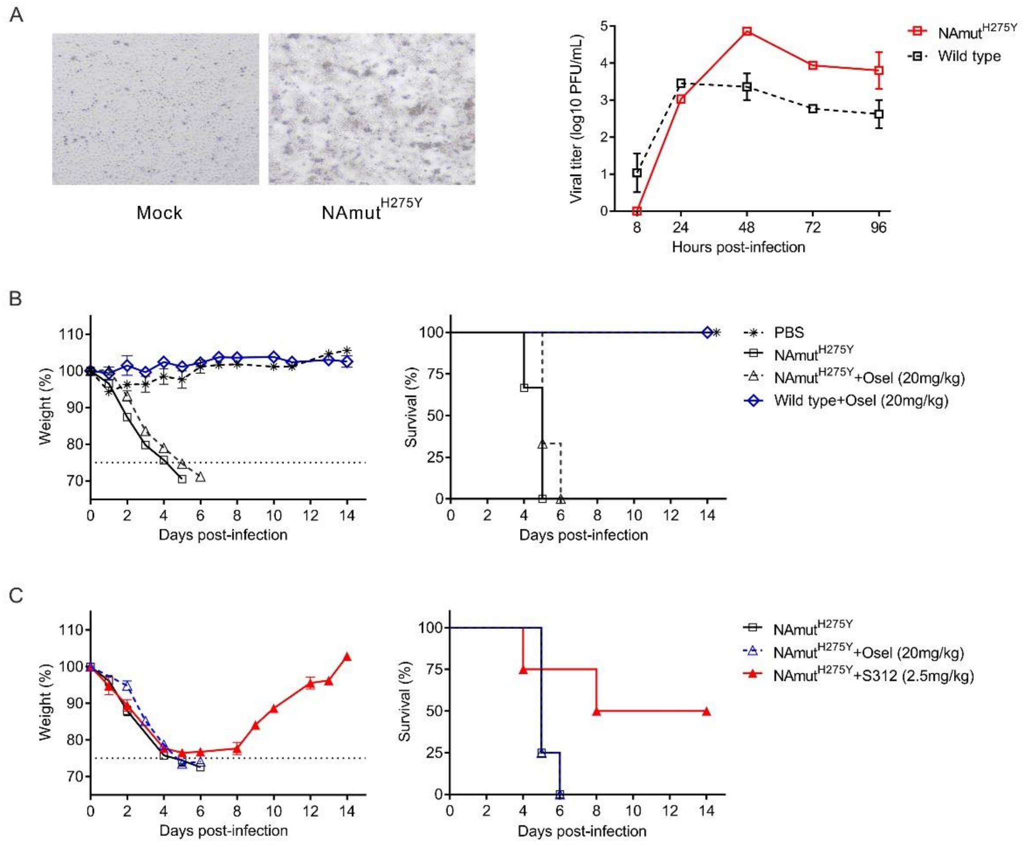


**Supplementary Fig. 2. S312 is active against Oseltamivir-resistant strain NAmut$^{H275Y}$. (A)** Viral growth curve. MDCK cells were infected with NAmut$^{H275Y}$ and influenza virus A/WSN/33 H1N1 (MOI=0.001). The supernatant of infected cells was collected at different hours post-infection. Viral titers were determined by plaque assay on MDCK cells. **(B, C)** BALB/c mice were intraperitoneally injected with PBS, Oseltamivir (20mg/kg) and S312 (2.5mg/kg) at 2.5 hours before infection, respectively. Then the mice intranasal infected with 5x10$^{4}$ PFU of NAmut$^{H275Y}$ and influenza virus A/WSN/33 H1N1. The mice were treated with the above compounds once a day. Bodyweight and survival of the mice were monitored for 14 days or until body weight reduced to 75%, respectively (n = 4 mice per group). And dotted line indicates endpoint for mortality (75% of initial weight). The results were presented as means ± SD.

**Supplementary Fig. 3**

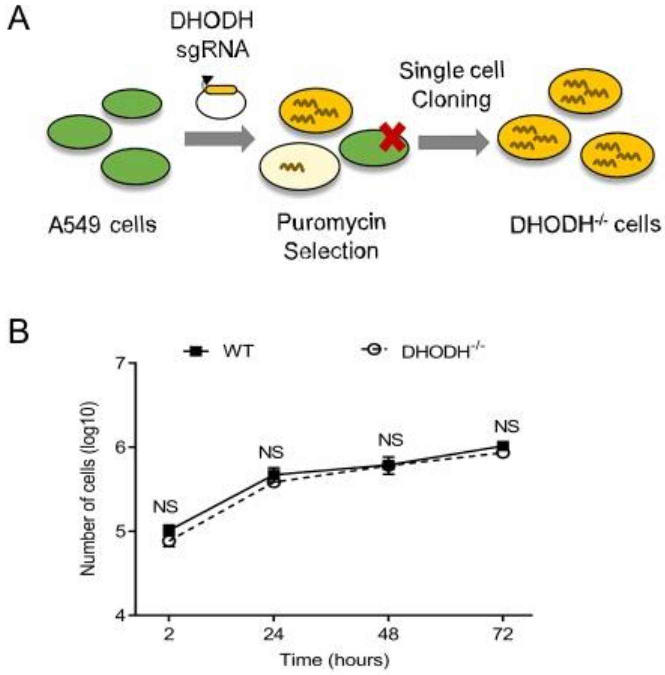


**Supplementary Fig. 3. Experimental procedure for DHODH knockout in A549 cells. (A)** Lentivirus-packaged sgRNA against DHODH was transduced into A549 cells and selected by puromycin with a concentration of 5 ug/ml. Subsequently, single-cell sorting was performed in a 96-well plate to grow single clones. The best KO cells with the lowest residue DHODH level were finally used. **(B)** Cell growth curve. A549 cells and DHODH$^{-/-}$ cells were respectively seeded in 48 well plates with $1\times10^5$ cells/well. And then counted by hemocytometer at each time points of 2h， 24h, 48h, and 72h. The results were presented as means ± SD. Statistical analysis, one-way ANOVA.

### Supplementary Fig. 4

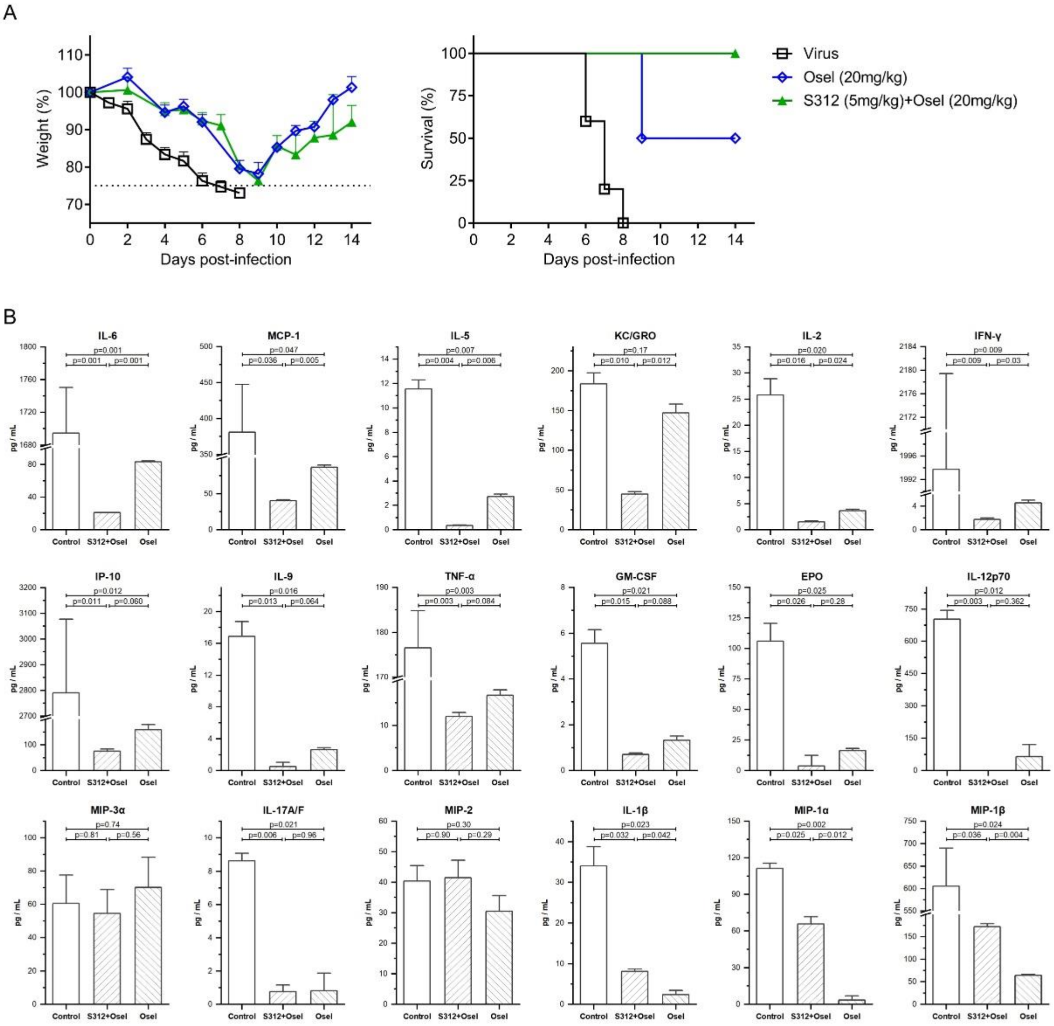


**Supplementary Fig. 4. Cytokine and chemokine measurements. (A)** BALB/c mice were intranasally infected with 2000 PFU of influenza virus A/WSN/33 H1N1. Then, give mice intraperitoneal injection (i.p.) with Oseltamivir (20mg/kg), S312 + Oseltamivir (5mg/kg + 20mg/kg) once a day. Bodyweight loss and survival of the mice were monitored for 14 days or until body weight reduced to 75%, respectively (n = 5 mice per group). And dotted line indicates endpoint for mortality (75% of initial weight). **(B)** The cytokines and chemokines were measured by Meso Scale Discovery (MSD). The data were expressed as mean ± SD and were used to create the bar charts with error bars. The statistical analyses were performed using one-way ANOVA followed by Turkey post-hoc test. The plot function, ANOVA and the post-hoc functions were provided by OriginPro 2020 SR1 (9.7.0.188). P<0.05 was considered statistically significant and therefore the "significance level" parameters of the above functions were set to 0.05.
